# CopR, a global regulator of transcription to maintain copper homeostasis in *Pyrococcus furiosus*

**DOI:** 10.1101/2020.08.14.251413

**Authors:** Felix Grünberger, Robert Reichelt, Ingrid Waege, Verena Ned, Korbinian Bronner, Marcell Kaljanac, Nina Weber, Zubeir El Ahmad, Lena Knauss, M. Gregor Madej, Christine Ziegler, Dina Grohmann, Winfried Hausner

## Abstract

Although copper is in many cases an essential micronutrient for cellular life, higher concentrations are toxic. Therefore, all living cells have developed strategies to maintain copper homeostasis. In this manuscript, we have analysed the transcriptome-wide response of *Pyrococcus furiosus* to increased copper concentrations and described the essential role of the putative copper-sensing metalloregulator CopR in the detoxification process.

To this end, we employed biochemical and biophysical methods to characterise the role of CopR. Additionally, a *copR* knockout strain revealed an amplified sensitivity in comparison to the parental strain towards increased copper levels, which designates an essential role of CopR for copper homeostasis. To learn more about the CopR-regulated gene network, we performed differential gene expression and ChIP-seq analysis under normal and 20 μM copper-shock conditions. By integrating the transcriptome and genome-wide binding data, we found that CopR binds to the upstream regions of many copper-induced genes. Negative-stain transmission electron microscopy and 2D class averaging revealed an octameric assembly formed from a tetramer of dimers for CopR, similar to published crystal structures from the Lrp family. In conclusion, we propose a model for CopR-regulated transcription and highlight the complex regulatory network that enables *Pyrococcus* to respond to increased copper concentrations.

## Introduction

The archaeal transcription system combines strategies and regulatory mechanisms known from eukaryotic as well as from bacterial species (1, 2). Archaea rely on a single RNA polymerase that synthesises all RNA species in the cell and is highly homologous to the eukaryotic RNA polymerase II (3). The presence of general transcription initiation factors (TATA box binding protein, Transcription factor B, Transcription factor E) and defined promotor elements (B recognition element, TATA box, initially melted region, initiation site) stresses the close relationship to eukaryotes especially for transcription initiation (4). In contrast, the fine-tuning process of gene expression is mainly achieved by bacterial-like transcriptional regulators (5). Positive or negative regulation is mediated by the binding of these transcription factors (TFs) to promoter regions of specific genes.

The genome of the hyperthermophilic euryarchaeon *Pyrococcus furiosus* contains a total number of 86 putative DNA-binding TFs. However, the exact function of most of these factors, which represent about 4 % of all open reading frames (ORFs), is unknown (6). In an attempt to close that knowledge gap, functional and structural aspects of some of these TFs have been analysed over the last two decades. While the regulation of sugar or sulfur metabolism and other changing environmental conditions have been studied in detail, the underlying mechanisms to maintain metal homeostasis are only poorly understood (7–11).

Playing an essential role in the cycling of elements, Archaea not only have to transform and make use of a variety of metals but also have to withstand elevated levels in the respective habitat (12). For many organisms, copper (Cu) is one of the essential trace elements used as a cofactor in a variety of proteins. These are mainly involved in electron transfers due to the ability to undergo redox changes from the reduced form Cu^+^ to the oxidised Cu^2+^. Despite its essential role, high intracellular concentrations are toxic for prokaryotic and eukaryotic cells. Copper catalyses the conversion of H2O2 to hydroxyl radicals via the Fenton reaction, which leads to oxidative damage of nucleic acids, proteins and lipids (13, 14). Cu^+^ also is a strong soft metal and can attack and destroy iron-sulfur proteins either by direct interaction or by blocking iron-sulfur cluster biogenesis (15, 16). To prevent cellular damage, all cells have developed various copper detoxification strategies. In prokaryotes, this is mainly achieved by active export of copper ions and in rarer cases by sequestration or exclusion (12, 17).

The two subfamilies of ATPases, P_1B-1_ (CopA) and P_1B-3_ (CopB), are the key players in cellular copper export. CopA transports Cu^+^, and CopB is proposed to transport Cu^2+^ (18, 19). To elucidate the mechanism of the exporting enzymes, the structures of homologous archaeal Cu-transporting ATPases CopA and CopB were studied in the hyperthermophilic *Archaeoglobus fulgidus.* The two enzymes seem to have different affinities for Cu^+^ and Cu^2+^ (20–22). Recent data, however, suggest that both subclasses, P_1B-1_ and P_1B-3_, have to be assigned as Cu^+^ transporters, which is consistent with the presence of only Cu^+^ in the reducing environment of the cytoplasm (23). Furthermore, a corresponding metallochaperone of the CopZ family is capable of reducing Cu^2+^ to Cu^+^ and is most likely involved in the transport of the reduced ion to CopA (24).

Many Archaea use the metallochaperone CopM, which contains a TRASH-instead of a heavy-metal-associated (HMA)-domain of the CopZ family (25). TRASH is a novel domain that has been proposed to be uniquely involved in metal-binding in sensors, transporters and trafficking proteins in prokaryotes (26). In addition to the specific binding of copper by chaperons, copper can also be buffered by small peptides like GSH and other reducing agents, to prevent cellular damage (27).

In several Archaea, the Cu-transporting ATPase and the copper chaperone are arranged in a conserved copper resistance gene cluster *(cop),* which also contains an additional gene, encoding for a DNA-binding transcriptional regulator. In previous studies, PF0739 has been bioinformatically predicted to be the copper-dependent regulator CopR in *P. furiosus* (25, 28, 29).

Based on biochemical data, *in vitro* analysis and growth experiments using knockout strains, CopR was proposed to play opposing regulatory roles in different Archaea: While in *Thermococcus onnurineus* the transcriptional regulator (TON_0836) represses *copA,* both transporter and chaperone are activated in *Saccharolobus solfataricus* (SSO2652) (28, 29).

Here, we have characterised the metal-sensing transcriptional regulator CopR in *P. furiosus.* First, we described the influence of different metal ions on the DNA-binding ability of CopR to the shared *copR/copA* promoter and analysed the growth of parental and *copR* knockout strains under increasing copper levels. We further performed a differential gene expression analysis (DGE) and chromatin immunoprecipitation with high-throughput sequencing (ChIP-seq) under normal and copper-shock conditions to elucidate the CopR-regulated gene network in *P. furiosus*. Integrating the genome-wide results with a more in-depth functional and structural characterisation, we propose that CopR acts as a dual regulator to maintain copper homeostasis.

## Material & Methods

### Strains, Plasmids, and Primers

All strains, plasmids and primers used in the study are listed in Supplementary Table 1.

### Construction of the *copR deletion strain*

For the construction of *P. furiosus* parental strain MURPf52 and Δ*copR* strain MURPf74 a modified genetic system was developed for *P. furiosus* DSM3638 allowing markerless disruption of genes onto the chromosome. This system is based on selection via agmatine-auxotrophy and counter selection via 6-methylpurine as described for *P. furiosus* COM1 strain and *T. kodakarensis* (30, 31).

First, for disruption of the *Pyrococcus pdaD* gene (PF1623; arginine decarboxylase gene) via a double-crossover event plasmid pMUR264 was constructed according to Kreuzer et al. (32). The first fusion PCR product containing upstream and downstream regions flanking the *Pf pdaD* gene encoding a Pyruvoyl-dependent arginine decarboxylase was created using the following two primer pairs: (Pf1622_fP_AscI/Pf1622_rP) and (Pf1624_fP/Pf1624_rP_NotI). The second fusion PCR product consisted of a two-gene resistance cassette which was needed for the selection-counter-selection system. The resistance cassette contained a *gdh* promoter, the *hmgCoA* reductase from *T. kodakarensis,* the region coding for the *xgprt* (PF1950, Xanthine-guanine phosphoribosyltransferase) and the histone A1 terminator sequence of *P. furiosus* (33). The first part was amplified using the primers: (SimV_NotI_F/SimV_Rv). For the second part, the primer pair: (Pf1624_fP_fus_2/Pf1624_rP_SbfI_N) was used. Both PCR products were combined with single-overlap extension PCR, ligated using a NotI restriction site and inserted into a modified pUC19 vector (32) using AscI and SbfI restriction sites.

However, all attempts to markerless delete the *Pf pdaD* gene using this construct were not successful and thus pMUR264 was modified to allow gene disruption via a single crossover event. To remove the second homologous downstream region, plasmid pMUR264 was amplified using the primer pair: (pUC19_SbfI_F/Pf1950_SbfI_R). The resulting PCR product was digested by SbfI and ligated. This plasmid was denoted as pMUR242 and used for transformation of *P. furiosus* as described (32, 33). To obtain the markerless double mutant MUR37Pf, circular plasmid DNA of pMUR242 and strain MURPf27 (32) were used and the corresponding transformants were selected with 10 μM simvastatin in SME-starch liquid medium supplemented with 8 mM agmatine sulfate at 85°C for 48 h. Pure cultures of the intermediate mutant MUR37Pf_i were obtained by plating the cells on solidified medium in the presence of 10 μM simvastatin and 8 mM agmatine sulfate. The integration of the plasmid into the genome by single cross-over was verified by analyzing corresponding PCR products.

Cultures of the correct intermediate mutant were washed with medium under anaerobic conditions to remove the simvastatin. In detail, 1.5 ml of a grown culture were centrifuged in an anaerobic chamber for 4 min at 6,000 g and resuspended in fresh culture medium without simvastatin. This procedure was repeated three times. For the counter selection, the cultures were grown in the presence of 50 μM 6-methylpurine and 8 mM agmatine sulfate to induce a second homologous recombination step to recycle the selection marker and to eliminate integrated plasmid sequences. Pure cultures were obtained by single cell isolation using an optical tweezer (34). Cultures had to be grown in the presence of 8 mM agmatine sulfate and 8 mM Inosine and Guanine (I+G). The genotype of the final mutant was confirmed by PCR and Southern blot experiments.

For markerless disruption of the *Pf copR* gene (PF0739), plasmid pMUR527 was constructed. First a modified resistance cassette had to be designed, which was needed for the selection-counter-selection system. The two-gene resistance cassette contained the *Pf pdaD* gene *including* the promoter and terminator region sequence of *P. furiosus* which was amplified using the primers: (PF1623F_Pr_BHI/PF1623R_Term). For the second part, the three primer pairs: (F_PF1950_P_F_Fu/R_PF1950_Prom, F_PF1950_Fs_P/PF1950_R and F_PF1950_Fus_T/PF1950_T_R_BHI) were used to amplify the promoter, coding and terminator region of the *Pf xgprt gene.* The four PCR products were combined with single-overlap extension PCR and subcloned into pUC19 vector via the SmaI restriction site. In the next step it was cloned via NotI and SbfI restriction sites into plasmid pMUR47 (32). The upstream and downstream flanking regions of the *Pf copR* gene were amplified using the primer pairs: (0739upAscIFW/0739up2RW and 0739dofus2FW/0739doNotIRW). Both PCR products were combined with single-overlap extension PCR and cloned into modified pMUR47 vector via the AscI and NotI restriction sites. The resulting construct (pMUR527) was verified by DNA sequencing.

Circular plasmid DNA and strain MURPf37 were used for transformation and selection was carried out in SME-starch liquid medium without agmatine sulfate and I+G at 85°C for 12 h. Pure cultures of the intermediate mutant MUR65Pf_i were obtained by plating the cells on solidified medium. The integration of the plasmid into the genome by single cross-over was verified by analyzing corresponding PCR products.

For the counter selection cells were plated on solidified medium containing 50 μM 6-methylpurine and 8 mM agmatine sulfate to induce a second homologous recombination step to recycle the selection marker and to eliminate integrated plasmid sequences. The genotype of the final mutant (MUR65Pf) was confirmed by PCR and cells had to be grown in the presence of 8 mM agmatine sulfate and 8 mM I+G.

To restore wild type growth properties (growth without agmatine sulfate and I+G) plasmid pMUR310 was created. The newly designed two-gene resistance cassette was amplified from the pUC19 subclone using the primer pair: (pYS_PF1623F_GA/pYS_PF1950R_GA). This PCR product was cloned into PCR-amplified (PF1623_pYSF_GA/PF1950_pYSR_GA) pYS3 plasmid (33) using NEB Gibson Assembly® Cloning Kit. Correctness of the construct was tested by Sanger sequencing. 1 μg of the circular plasmid was transformed into MURPf37 and MURPf65 as described (32, 33). Selection was carried out in 1/2 SME liquid medium without agmatine sulfate and I+G at 85°C for 12 h. Pure cultures of the mutant MUR52Pf and MURPf74 were obtained by plating the cells on solidified medium. Plasmid stability was verified by re-transformation into *E. coli* and DNA sequencing of purified plasmids. Final mutants could be grown without agmatine sulfate and I+G supplementation.

RNAP, TBP, TFB: For *in vitro* transcription assays and EMSA analysis, we used RNAP purified from *P. furiosus* cells and recombinant TBP and TFB as described previously (33, 35, 36).

### CopR, CopRΔTRASH, CopRΔHIS and CopRΔHISΔTRASH

#### Cloning and expression

The gene sequence of PF0739 was amplified from genomic DNA of *P. furiosus* using primers with additional BamHI and NdeI restriction recognition sites. The PCR product was cloned into vector pET-30b (NEB) using the respective restriction sites. PF0739 protein variants lacking potential metal-sensing domains (ΔTRASH, ΔHIS, ΔHISΔTRASH) were based on the full-length plasmid version and ligated after amplification using one phosphorylated primer, respectively (see Supplementary Table 1). Subsequently, the constructs were transformed into *E. coli* DH5-α for amplification and grown on Kanamycin (50 μg /ml) supplemented LB media. Next, the constructs were transformed into *E. coli* BL21 STAR™ (DE3) expression strain and grown on Kanamycin (50 μg /ml) supplemented LB medium at 37°C. Protein expression was induced by addition of 0.5 mM IPTG to the cell culture medium at an OD_600_ of about 0.6. Cultures were further cultivated at 18°C overnight, before harvesting the cells by centrifugation at 10,000 g for 10 min at 4°C. Cells were stored at −80°C until protein purification.

#### Cell disruption and pre-purification

For the purification of PF0739 and PF0739 variants, cells were first resuspended in 50 ml low salt buffer containing 40 mM HEPES (pH 7.5), 80 mM ammonium sulfate, 1 mM EDTA, 10% glycerol (w/v) and a protease inhibitor tablet (Roche). The cell lysis was done by sonification on ice, whereby breakage efficiency was monitored at a light microscope. After cell disruption, DNase I (Roche) was added and all cultures incubated at 37°C for 1 hour. In the next step, the lysate was centrifuged at 48,000 g for 20 min at 4°C and the supernatant was transferred to a new tube. A pre-purification step was carried out by applying a heat treatment of 90°C for 15 min and subsequent centrifugation at 48,000 g for 20 min at 4°C.

#### Affinity and size exclusion chromatography

The supernatant containing the protein of interest was filtered and loaded onto a 5-ml HiTrap™ Heparin HP column equilibrated with low salt buffer. Next, the protein was eluted by gradually increasing the buffer concentration of the high salt buffer (compare low salt, but 1 M ammonium sulfate). Fractions containing PF0739 (checked on SDS-PAGE) were further purified using size exclusion chromatography by pooling and concentrating of the relevant fractions and loading onto a 24-ml HiLoad™ 10/300 GL Superdex™ 200 column pre-equilibrated with low salt buffer. This column was also used to study multimerization of PF0739 (data not shown).

Growth experiments using an optical device: *P. furiosus* was cultivated under anaerobic conditions in 40 ml ½ SME medium supplemented with 0.1 % yeast extract, 0.1 % peptone and 40 mM pyruvate at 95°C, as described previously (33, 37). For growth comparison experiments, the medium was supplemented with different CuSO_4_ concentrations (compare Fig. 2) and each condition for MURPf52 (parental strain) and MURPf74 *(ΔcopR* strain) was recorded in biological triplicates during 48 hours of incubation by measuring the turbidity changes *in situ* using a photodiode and a LED with 850 nm as light source. The recorded values were converted to cell/ml by using a calibration curve with known cell concentrations, calculated in a Thoma counting chamber (0.02-mm depth; Marienfeld, Lauda-Königshofen, Germany) using phase-contrast microscopy.

### Differential gene expression analysis

#### Growth conditions and RNA isolation

*P. furiosus* parental strain MURPf52 was grown in standard medium at 95°C to late-exponential phase. After reaching a cell density of 1×10^8^, cells were either shocked by adding 20 μM CuSO_4_ (copper-shock) or left untreated (control) and incubated for 30 minutes. The experiment was performed in biological triplicates. Total RNA was isolated using the Monarch RNA purification Kit (NEB), including the recommended genomic DNA removal by on-column DNase treatment, according to the instructions of the manufacturer. Quantity, quality and integrity were measured using Nanodrop One, Qubit RNA HS assay kit (Thermo Fisher Scientific) and the Prokaryote total RNA Nano Kit on a Bioanalyzer to measure RIN values (Agilent).

#### Library preparation and sequencing

Library preparation and RNA-seq were carried out as described in the Illumina TruSeq Stranded mRNA Sample Preparation Guide, the Illumina HiSeq 1000 System User Guide (Illumina, Inc., San Diego, CA, USA), and the KAPA Library Quantification Kit - Illumina/ABI Prism User Guide (Kapa Biosystems, Inc., Woburn, MA, USA). In brief, 100 ng of total RNA from *P. furiosus* was fragmented to an average insert size of 200-400 bases using divalent cations under elevated temperature (94°C for 4 minutes), omitting the mRNA purification step with poly-T oligo-attached magnetic beads. Next, the cleaved RNA fragments were reverse transcribed into first strand cDNA using reverse transcriptase and random hexamer primers. Actinomycin D was added to improve strand specificity by preventing spurious DNA-dependent synthesis. Blunt-ended second strand cDNA was synthesized using DNA Polymerase I, RNase H and dUTP nucleotides. The incorporation of dUTP, in place of dTTP, quenched the second strand synthesis during the later PCR amplification, because the polymerase does not incorporate past this nucleotide. The resulting cDNA fragments were adenylated at the 3’ ends, the indexing adapters were ligated, and subsequently specific cDNA libraries were created by PCR enrichment. The libraries were quantified using the KAPA SYBR FAST ABI Prism Library Quantification Kit. Equimolar amounts of each library were used for cluster generation on the cBot with the Illumina TruSeq SR Cluster Kit v3. The sequencing run was performed on a HiSeq 1000 instrument using the indexed, 50 cycles single-read (SR) protocol and the TruSeq SBS v3 Reagents according to the Illumina HiSeq 1000 System User Guide. Image analysis and base calling resulted in .bcl files, which were converted into FASTQ files with the bcl2fastq v2.18 software. Library preparation and RNA-seq were performed at the service facility “KFB - Center of Excellence for Fluorescent Bioanalytics” (Regensburg, Germany; www.kfb-regensburg.de).

#### Data analysis using the DESeq2 pipeline

For differential gene expression analysis, rRNA-derived Illumina reads were first removed using SortmeRNA (38). Next, reads in FASTQ format were quality/length/adapter trimmed using trimmomatic (v. 0.36) in singleend-mode (39). Therefore, we allowed for a minimum length of 12 bases and a cut-off Phred score of 20, calculated in a sliding window of 4 bases. Wee used the STAR aligner (v. 2.5.4) to map the reads to a recently published updated version of the *P. furiosus* genome (40, 41). Mapping statistics are included in the Supplementary Table 2. The sorted BAM files were then used to generate count tables using featureCounts (42). Differential gene expression analysis was performed using the DESeq2 pipeline (43). Furthermore, we used the *apeglm* method for effect size shrinkage and calculation of fold changes (44). All steps of the analysis, including the generation of plots were performed using R and can be found at www. github.com/felixgrunberger/CopR (45). Enrichment analysis of archaeal cluster of orthologous genes (arCOGs) was performed by extracting the gene specific arCOG information from the arCOG database (ftp://ftp.ncbi.nih.gov/pub/wolf/COGs/arCOG/) and calculation using the goseq package, which allowed for custom genome sets (46, 47).

#### Confirmation of data using RT-qPCR

RT-qPCR reactions were performed similar as described previously (36). In short, total isolated RNA was reverse transcribed using the ProtoScript® II First Strand cDNA Synthesis Kit (NEB), according to the manufacturer’s instructions and using a random primer mix (Promega). The reactions were assembled in triplicates using the qPCRBio SyGreen Mix Lo-Rox Kit (PCRBiosystems) with reverse transcribed cDNA from the first step in a 1:10 dilution, including a control reaction that lacked the reverse transcriptase (-RT) and a no template control (NTC). RT-qPCR reactions were run on a Rotor-Gene Q cycler (Qiagen) in a three-step protocol: 95°C - 10’ for one cycle; 95°C – 30”, 58°C – 30”, 72°C – 30” for 40 cycles. Data evaluation was done using the corresponding Rotor-Gene Q software package (Qiagen). Relative expression levels were calculated using the delta-delta Ct method (2^-ΔΔCt^), by comparing the Ct values from biological triplicates of the gene of interest to a house-keeping gene *pf0256.* The applicability of *pf0256* as a calibrator was evaluated before (36).

### ChIP-seq analysis

#### Immunoprecipitation

We used an adaption of a ChIP-seq protocol that was established previously for P. furiosus (48). P. furiosus cells were grown under anaerobic conditions in serum bottles containing 40 ml ½ SME medium at 95°C as described earlier. After the cells reached a density of 2×10^8^, formaldehyde was injected into the flask to a final concentration of 0.1 % (v/v). After 60 seconds the crosslinking reaction was stopped by addition of glycine to a final concentration of 15 mM (v/v). For the copper-treated samples, the cells were shocked with 20 μM CuSO_4_ for five minutes before the crosslink reaction was induced. Cell disruption and DNA fragmentation was performed in one step via sonication for 25 minutes using the ultrasonic homogenizer Sonopuls HD 2070 (Bandelin, Berlin, Germany) until an average fragment length of 250 to 400 bp. The insoluble particles were removed by centrifugation. For determination of the DNA concentration and fragment length, 1 volume of crude cell extract was mixed with 4 volumes of ChIP elution buffer (10 mM Tris, pH 8.0, 1% (w/v) SDS, 0.1 mM EGTA) and incubated over night at 65°C. After RNase treatment, the DNA was purified using the NucleoSpin® Gel and PCR Clean-up Kit (Macherey-Nagel). The DNA concentration was determined using the Qubit dsDNA BR Assay Kit (Thermo Fisher Scientific) and the fragment length by agarose gel electrophoresis. For immunoprecipitation (IP) 100 μl Protein G beads (Dynabeads, Invitrogen) were coupled to 3920 μg serum antibodies against CopR (PF0739), according to manufacturer’s instructions. Polyclonal antibodies were produced by Davids Biotechnology (Regensburg, Germany) from recombinantly expressed and purified CopR. 100 μl of antibody-beads complexes were mixed with 900 μl of P. furiosus crude extract adjusted to a DNA concentration of 4.44 ng/μl (4 μg DNA/sample) in PBST. The samples were incubated with rotation for two hours at room temperature. The immunoprecipitated samples were placed on a magnet, the supernatant was discarded, and the bead-pellet was washed 2x with low salt buffer (50 mM HEPES, pH 7.4, 150 mM NaCl, 1 mM EDTA, 0.1% (w/v) SDS, 0.1% (w/v) Deoxycholic acid, 1% (v/v) Triton X-100), 1x with high salt buffer (50 mM HEPES, pH 7.4, 500 mM NaCl, 1 mM EDTA, 0.1% (w/v) SDS, 0.1% (w/v) Deoxycholic acid, 1% (v/v) Triton X-100), 1x with ChIP wash buffer (10 mM Tris, pH 8.0, 250 mM LiCl, 1 mM EDTA, 0.5% (v/v) Nonidet P-40, 0.5% (w/v) Deoxycholic acid) and 1x with TE buffer (10 mM Tris, pH 8.0, 0.1 mM EDTA, 20 % (v/v) Methanol, 25 mM Tris, 192 mM Glycine) (49). Each washing step was done with 1 ml buffer by rotation for one minute. To elute the immuno-bound DNA from the beads, the bead-pellet was resuspended in 25 μl ChIP elution buffer, transferred to a PCR cup and incubated for 10 minutes at 65°C. The cup was placed on a magnet, the supernatant was transferred to a new cup, the bead-pellet was resuspended in 25 μl TE buffer supplemented with 0.67% SDS (v/v) and incubated for 10 minutes at room temperature. Afterwards, both eluates were combined in one PCR cup. For the input sample, 400 ng DNA of P. furiosus crude extract was mixed 1:4 with ChIP elution buffer. Eluted complexes and input samples were incubated overnight at 65°C to reverse the crosslink. After the incubation, the samples were treated with RNase A (0.1 mg/ml final concentration) for 15 minutes at 37°C and Proteinase K (0,2 mg/ml final concentration) for 15 minutes at 65°C. ChIP-DNA and input DNA were purified using the NucleoSpin® Gel and PCR Clean-up Kit (Macherey-Nagel). The DNA concentration of the input DNA was determined using the Qubit dsDNA BR Assay Kit (Thermo Fisher Scientific).

#### Library preparation and sequencing

Library preparations were done using the NEBNext® UltraTM II DNA Library Prep Kit for Illumina® with the NEBNext® Multiplex Oligos for Illumina® Index Primers Set 2 and 3 according to the manufacturer’s protocol. Library quantification was done with the NEBNext® Library Quant Kit for Illumina® according to manufacturer’s instructions. Before sequencing, the libraries were pooled in equimolar ratios. The library pool was quantified with the KAPA SYBR FAST ABI Prism Library Quantification Kit (Kapa Biosystems, Inc., Woburn, MA, USA) and used for cluster generation on the cBot with the Illumina TruSeq SR Cluster Kit v3. Sequencing was performed on a HiSeq 1000 instrument controlled by the HiSeq Control Software (HCS) 2.2.38, using the indexed, 50 cycles single-read (SR) protocol and the TruSeq SBS v3 Reagents according to the Illumina HiSeq 1000 System User Guide. Image analysis and base calling were done by the Real Time Analysis Software (RTA) 1.18.61. The resulting .bcl files were converted into FASTQ files with the CASAVA Software 1.8.2. Sequencing was performed at the service facility “KFB - Center of Excellence for Fluorescent Bioanalytics” (Regensburg, Germany).

#### Analysis of ChIP-seq data

FASTQ files were quality/length/adapter trimmed with trimmomatic (v. 0.36) in single-end-mode using a minimum-length of 40 bp, a cut-off Phred score of 20 (39). Reads were mapped to the P. furiosus genome using Bowtie 2 (v. 2.2.3) with default settings (50). SAM files were converted to sorted BAM files using samtools and extended towards the 3’direction by their fragment-size to better represent the precise protein-DNA interaction (51, 52). Position-specific enrichments were calculated by extracting the mean counts of biological triplicates from bed files for the ChIP-sample, comparison to the input files and taking the log_2_.

#### Confirmation of data using quantitative real-time PCR (RT-qPCR)

qPCR primer pairs were designed using the Primer3 software package and quality assessed. qPCR reactions were assembled as technical triplicates using 2x qPCRBioSyGreen Mix separate Rox kit in a total volume of 10 μl. Primers were added to a final concentration of 0.3 μM. 6 μl of the master mix were mixed with 4 μl of template DNA or H_2_O_DEPC_ as NTC. The specificity of the PCR product was verified by melting curve analysis. qPCR reactions were run on the Rotor Gene Q cycler with a three-step PCR program described in the RNA-seq section. Replicates with a deviation >0.5 were excluded from the analysis. The fold enrichment was again calculated according to the delta-delta Ct method.

### In vitro assays

#### Electrophoretic mobility shift assay (EMSA)

DNA templates were obtained from genomic DNA by PCR amplification with the corresponding primer pairs (see Supplementary Table 1). One of the two primers was labelled at the 5’-end with a fluorescent dye. 20 nM DNA was assembled in a 15 μl reaction volume containing: 50 ng competitor DNA (Hind-III-digested λ DNA), 670 μM DTT, 20 μg/ml BSA, 6.7 % glycerol, 40 mM HEPES (pH 7.4), 80 mM (NH4)2SO4 and various amounts of proteins and metals, as described in the results part. The reactions were incubated for 5 minutes at 70°C and analysed using a non-denaturing 5 % polyacrylamide gel. After electrophoresis, the DNA fragments were visualized with a Fujifilm FLA-5000 fluorescence imager.

#### DNase I footprinting

DNase I footprinting was performed as previously described (35). In short, the DNA template containing the promoter regions of *pf0739* and *pf0740* was obtained from genomic DNA by PCR amplification (see Supplementary Table 1). HEX-labelled primers were used in two separate reactions for strand-specific labelling. 4.4 nM template DNA was assembled in a 15 μl reaction volume containing: 40 mM Na-HEPES (pH 7.5), 125 mM NaCl, 0.1 mM EDTA, 0.1 mg/ml BSA, 1 mM DTT, 0.5 mM MgCl_2_ and 2.8 μM CopR, 1 μM TBP and 0.8 μM TFB according to Fig. 5. After incubation for 20 minutes at 70°C, 0.05 units of DNase I (Fermentas) was added and incubated for another minute at 70°C. The reaction was terminated by the addition of 5 μl 95 % formamide and incubation for 3 minutes at 95°C. The DNA was precipitated with ethanol and resuspended in 2-4 μl formamide buffer. A DNA sequencing ladder was generated using a DNA cycle Sequencing Kit (Jena Bioscience) according to manufacturer’s instructions. Samples were loaded onto a 4.5 % denaturing polyacrylamide gel and analysed using an ABI 377 DNA sequencer.

#### In vitro transcription assay

*In vitro* transcription assays were performed similar as described previously (35, 36). To this end, template DNAs containing the promoter regions of the respective gene were amplified from genomic DNA or plasmid pUC19/*gdh* by PCR amplification (see Supplementary Table 1). The reactions were assembled in a total volume of 25 μl containing: 2.5 nM template, 5 nM RNAP, 30 nM TFB, 95 nM TBP, 40 mM Na-HEPES (pH 7.4), 250 mM KCl, 2.5 mM MgCl2, 5 % (v/v) glycerol, 0.1 mM EDTA, 0.1 mg/ml BSA, 40 μM GTP, 40 μM ATP, 40 μM CTP, 2 μM UTP, and 0.15 MBq (110 TBq/mmol) [α-32P]-UTP, if not indicated otherwise (see Fig. 5). After incubation for 10 minutes at 80°C, the RNA transcripts were extracted by phenol/chloroform, denatured in formamide buffer for 3 minutes at 95°C and separated on a denaturing 6 % polyacrylamide gel. Finally, the transcripts were visualised using a fluorescence imager (FLA-5000, Fuji, Japan).

### Negative-stain transmission electron microscopy and image analysis by 2D class averaging

For transmission electron microscopy (TEM), protein solutions with a concentration of 220 ng/μl were chosen. Samples were negatively stained with a solution containing 2 % (w/v) uranyl acetate (UAc) in presence of 0.005 % n-dodecyl β-D-Maltopyranosid (DDM). A carbon film coated grid - 400 Square Mesh (Plano GmbH), was incubated with 3 μL of protein sample for 45 seconds. Excess stain was blotted off using filter paper and samples washed using 3 μl of a 2 % UAc solution. The blotting and washing procedures were repeated and the sample finally air-dried and stored at room temperature. Negative-stained grids were imaged on a TEM JEOL-2100F (200 kV) equipped with a 4k x 4k F416 camera with CMOS chip/detector, TVIPS at a 50K magnification (0.211 nm/pixel) with a defocus range from −0.5 to −1.4 μm. The contrast transfer function for a total of 32 micrographs was determined with CTFfind4 (53). Subsequently, the particles were extracted with a mask of 180 Å and processed in RELION 3.0 (54) to yield the 2D class averages.

## Results

### Pyrococcus copR is part of the conserved archaeal cop gene cluster

A conserved *cop* resistance gene cluster plays a critical role in copper homeostasis in Archaea. The gene cluster has been identified in various archaeal species using comparative genomics (25). This *cop* cluster consists of a copper-exporting PiB-ATPase CopA *(Ferroplasma acidarmanus* Fer1: CopB), a transcriptional regulator CopR *(Saccharolobus solfataricus* P2: CopT, *F. acidarmanus* Fer1: CopY) and occasionally the metallochaperone CopT (*S. solfataricus* P2: CopM, *F. acidarmanus* Fer1: CopZ) (Fig. 1A) (25, 28, 55, 56).

**Figure 1:**
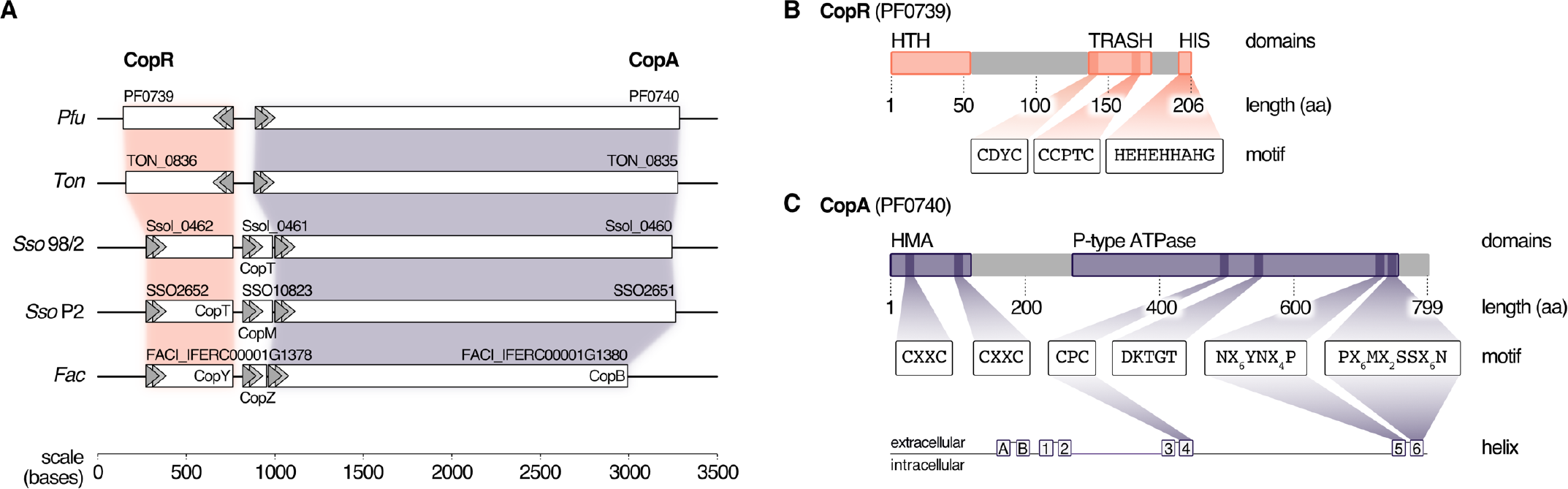
PF0739 (CopR) is part of the conserved archaeal cop cluster in *Pyrococcus furiosus*. **A**, Copper regulation in Archaea is achieved by a highly conserved cop gene cluster consisting of a transcriptional regulator (CopR/CopT/CopY), a transporter (CopA/CopB) and optionally a chaperone (CopT/CopM/CopZ). The organisation of the cop cluster in *P. furiosus* was compared to cop clusters of other archaeal organisms (*Ton = Thermococcus onnurineus, Sso96/2 = Saccharolobus solfataricus 96/2, SsoP2 = Saccharolobus solfataricus P2, Fac = Ferroplasma acidarmanus*) (25, 28, 29, 55). Genes are drawn to scale; directionality is indicated by arrows. **B**, Schematic representation of the transcriptional regulator (CopR) encoded by *pf0739* in *P. furiosus*. The regulator consists of a N-terminal helix-turn-helix (HTH) domain that mediates DNA binding, and a metal-sensing TRASH domain (26). The C-terminal Histidine-rich sequence (HIS) is only found in *Thermococcales*. Further sequence features of the domains are highlighted. **C**, Bioinformatical analysis of conserved amino acids in transmembrane helices 4, 5 and 6 classify PF0740 as the copper exporter CopA.

In *Pyrococcus furiosus,* the transcriptional regulator CopR is encoded by the gene *pf0739,* which is in divergent orientation to *copA* (*pf0740*). It consists of an N-terminal helix-turn-helix domain and a C-terminal metal-sensing TRASH domain together with a Histidine-rich region (HIS), the latter one is only present in the order *Thermococcales* (Fig. 1B) (26). To unravel more details about the function of CopR in *P. furiosus,* we expressed the wild type protein together with three mutants in *Escherichia coli* and tested the proteins for DNA binding using gel-shift assays. Here, the intergenic region between *copR* and *copA* served as target DNA. Binding of CopR to the target region with increasing protein concentrations resulted in a specific protein-DNA shift (Supplementary Fig. 1). The mutated variants, lacking the putative metal-binding domains, TRASH, HIS or both, showed a very similar DNA binding affinity for all variants, indicating that DNA binding is primarily mediated by the HTH domain whereas the metalbinding domains are dispensable for DNA binding.

To determine both selectivity and sensitivity towards the recognized metal of the CopR/CopA system in *P. furiosus,* we performed two types of experiments: First, motif analysis of the heavy metal-binding domains (HMBDs) classified PF0740 as a copper-exporting ATPase of type 1B (Fig. 1C) (57, 58). Secondly, increasing concentrations of different metal ions (AgNO3, CuSO4, FeCl3 and, CoCl2) were supplemented in the binding assays of CopR. All ions reduced the binding affinity, but the most potent effect was observed for copper and silver ions (Supplementary Fig. 2A). In contrast to the copper-induced release in the EMSA analysis using the full-length CopR, the effect was slightly reduced in the case of a CopRΔHIS mutant and significantly reduced for a CopRΔTRASH mutant (Supplementary Fig. 2B,C,D). It is interesting to note that lower metal concentrations resulted in a smear with reduced mobility and higher levels in an increasing amount of released DNA. The minimal concentration used in the binding assays that resulted in CopR release was 12.5 μM CuSO4, which is very similar to the detection range of CopR in *T. onnurineus* (79 % amino acid sequence identity) (28). Taken together, these results indicate that the CopR/CopA system is involved in copper regulation in *P. furiosus*.

### Deletion of copR transcriptional regulator leads to a copper-sensitive phenotype

To learn about the importance of CopR for copper-detoxification in *P. furiosus*, a *copR* deletion mutant was constructed, using an established genetic system in this hyperthermophilic organism (32, 33). Growth analysis of this deletion mutant (MURPf74) in comparison to the parental strain (MURPf52) was performed using increasing amounts of copper (0 to 100 μM). Starting at sub-lethal concentrations, we observed prolonged lag phases and reduced cell densities in both strains (Fig. 2).

**Figure 2:**
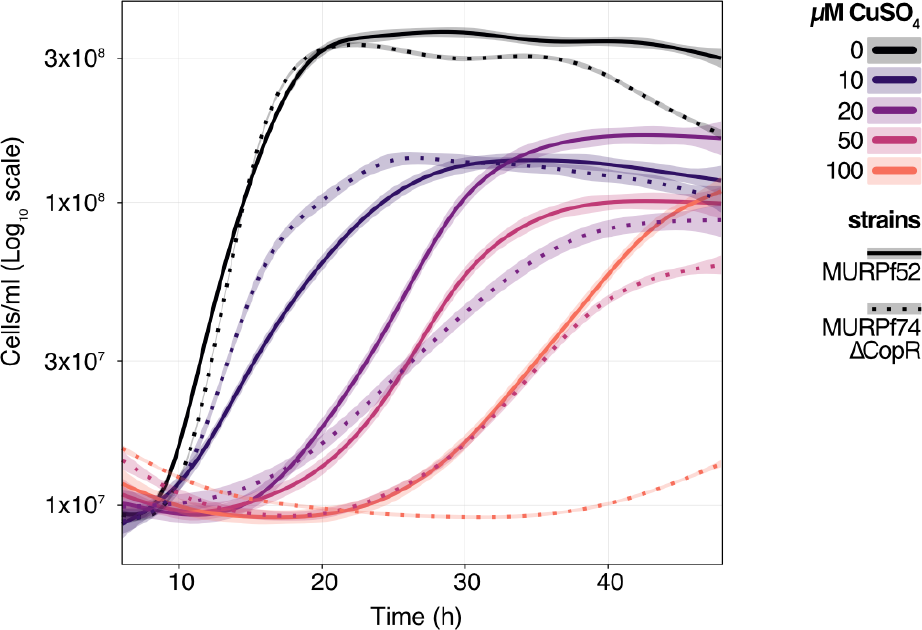
Growth analysis of the *P. furiosus* parental strain (MURPf52) and CopR-knockout strain (MURPf74) in the presence of CuSO_4_. Triplicates of 40 ml cultures were analysed in standard medium at 95°C, supplemented with 0 to 100 μM CuSO_4_ (indicated by colour scale). Growth was recorded during 48 hours of cultivation of the parental strain (solid lanes) and the *copR* deletion strain (dashed lines). Each curve represents the fitted line of three independent experiment, with the shaded area displaying the confidence interval (0.99).

While the collected curves were almost identical for 0 and 10 μM CuSO_4_, higher copper concentrations caused a significant effect on the growth of the knockout strain. In contrast to the parental strain, the growth of the *copR*-disrupted strain was almost completely abolished in the presence of 100 μM copper. This finding supports the idea that copper homeostasis is a tightly controlled system with a sensitivity in the μM-range and indicates that CopR acts as a transcriptional activator of the copper-exporting ATPase *pf0740* in *P. furiosus.*

### Characteristics of the P. furiosus transcriptome in response to a copper-shock

To investigate the role of CopR in the copper regulation network in *P. furiosus*, we applied an integrative approach, combining differential gene expression (DGE) analysis and genome-wide binding analysis by ChIP-seq. For DGE, we cultivated the parental strain (MURPf52) until the middle of the log phase, shocked the cells with 20 μM CuSO_4_ for 20 minutes and isolated the RNA for next-generation sequencing. PCA analysis confirmed that indeed the copper-shock (and not handling of the biological replicates) caused most of the variance in the experimental setup (Fig. 3A). Hence, we were able to compare the transcriptomic changes primarily due to copper shock and not as a result of a general stress response or cell death. By analysing the transcript abundances, we could confirm the essential role of *copA* in removing excess ions from the cell, observing a 70-fold up-regulation of the mRNA levels after copper treatment (Fig. 3B). Altogether, 34 genes were up-regulated, but not a single gene was significantly down-regulated by more than 2-fold (Fig. 3C,D). RT-qPCR experiments verified the up-regulation of two of the most prominent genes (PF0740, PF0738.1n) (Supplementary Fig. 3). Notably, *pf0727* is among the most up-regulated genes (105-fold). Based on the domain annotation and strong induction upon copper treatment, PF0727 is most likely the missing chaperone in the *cop* cluster in *P. furiosus.* Due to the presence of an HMA domain instead of a TRASH domain the protein belongs to the CopZ and not to the CopM family. A closer look at the clusters of archaeal orthologous genes (arCOGs) revealed that most of the up-regulated genes belong to the groups O (posttranslational modification, protein turnover, chaperones), S (function unknown) and P (inorganic ion transport and metabolism) (Fig. 3E,F) (47). The group of the hypothetical genes consists of eight candidates, including *pf0738.1n,* which exhibits the most substantial up-regulation (290-fold).

**Figure 3:**
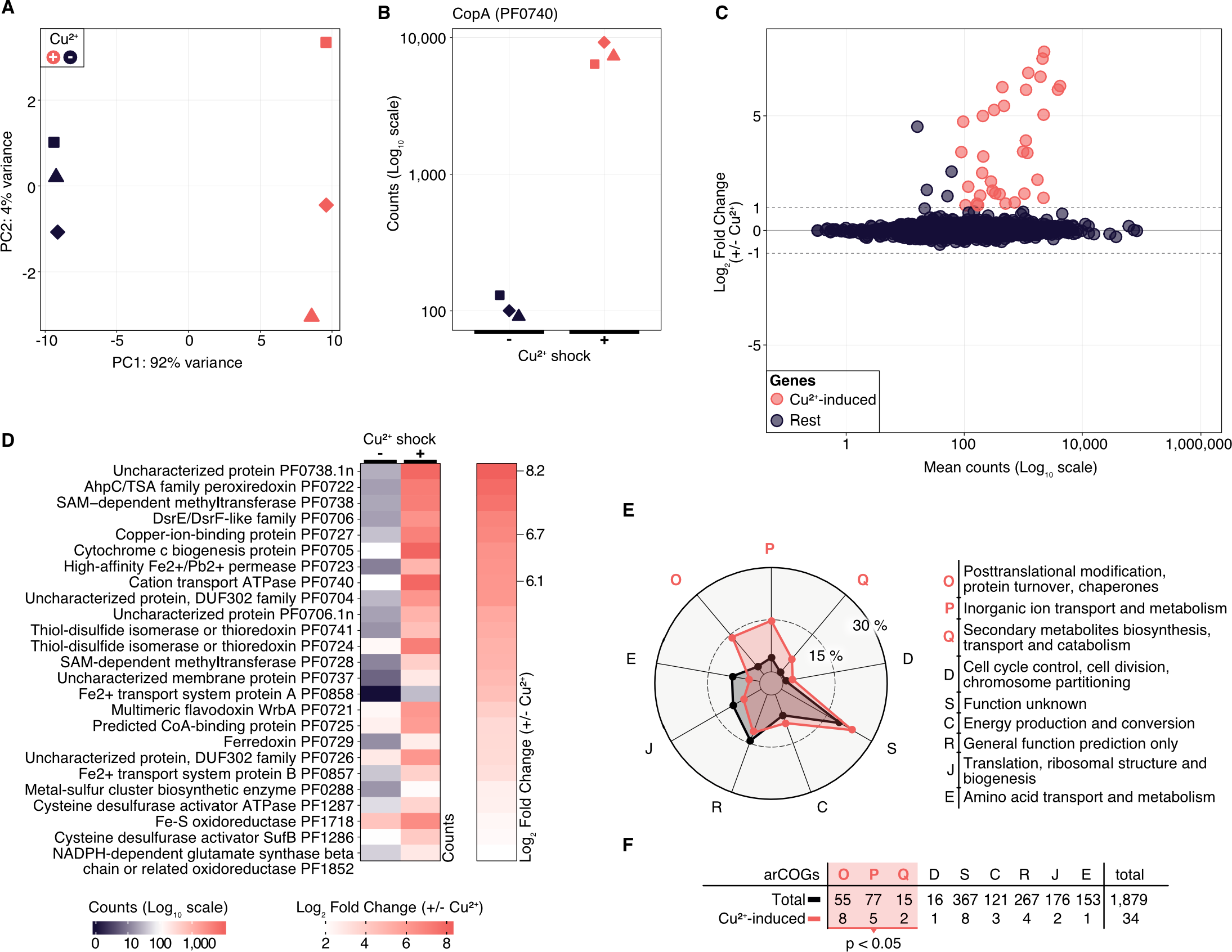
Differential gene expression analysis of *P. furiosus* after 20 min copper shock with 20 μM CuSO_4_. **A**, Principal component analysis of variance stabilized transformed RNA-Seq read-counts of normal conditions (dark blue) and Cu^2+^ shock conditions (red) shows that most of the variance in the experimental setup is caused by the treatment of the cells. Replicates are indicated by different shapes. **B**, Comparison of raw RNA-Seq readcounts mapping to PF0740 shows a 6.14 Log_2_ Fold Change after Cu^2+^ shock (adjusted p value = 1.80e-173). **C**, MA plot showing the distribution of mean read counts (log_10_ scale) against log_2_ fold change. 34 genes that are more than 2-fold up-regulated with an adjusted p-value <0.05 are highlighted in red. **D**, Heatmaps for mean read-counts for control and Cu^2+^ shock condition are shown for the 25 most significant regulated genes. **E**, Enrichment analysis of archaeal clusters of orthologous genes (arCOGs) found in the 34 significantly up-regulated genes (compare panel c). Contribution of each category is calculated in percentage and compared to the total background set with significantly up-regulated categories marked in orange. **F**, Enrichment analysis is based on the comparison of the total number of genes found in an arCOG category (Total) and the number of genes that are significantly up-regulated (>2 fold).

### Integrative RNA-seq and ChIP-seq identifies CopR targets

It is interesting to note that the 14 most up-regulated genes are located within a 28 kb region of the genome. To answer the question if the transcriptional activation of these genes upon intoxication is connected to the binding of CopR, we performed a ChIP-seq experiment with and without copper shock (20 μM CuSO_4_). The results from these experiments demonstrated a very similar CopR occupancy independent of the copper treatment (Fig. 4A). This finding is in agreement with the EMSA analysis, where a considerable amount of the transcriptional regulator remained bound to the DNA after addition of 25 μM CuSO_4_ (comparable amount as used in the *in vivo* experiments, see Supplementary Fig. 2). Furthermore, the binding pattern of CopR overlaps with the upstream regions of up-regulated genes or operons under both conditions, which confirms specific binding of CopR, as well as the possible role in transcriptional activation.

**Figure 4:**
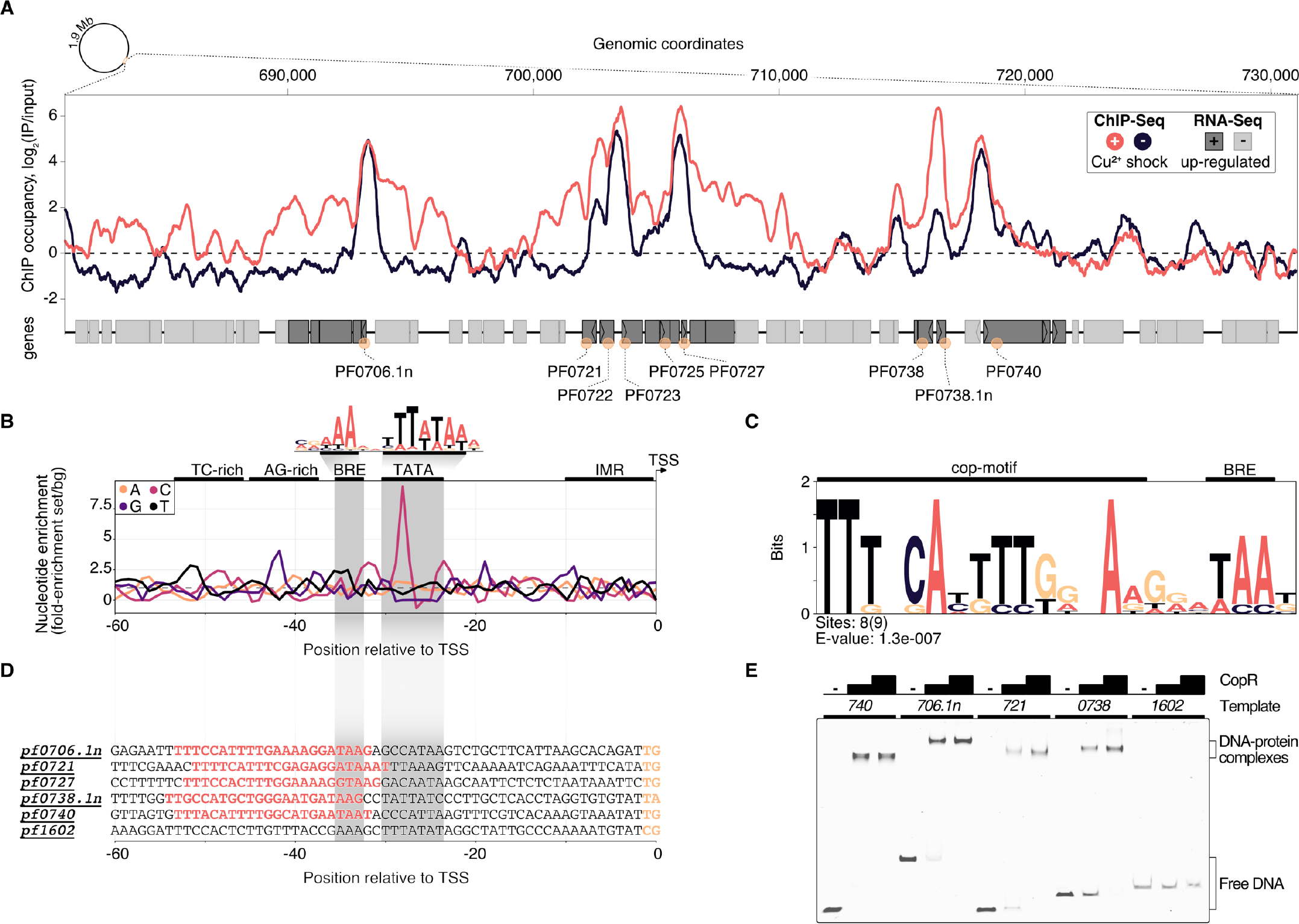
ChIP-seq and integration with differential gene expression data identifies CopR as a global regulator of copper homeostasis in *P. furiosus.* **A**, ChIP occupancy of CopR zoomed to genomic region ~680,000 to 730,000, which contains the 14 most up-regulated genes from the differential gene expression analysis (compare Fig. 3). ChIP-seq curves were generated for Cu^2+^ shocked (red) and untreated (dark blue) samples by comparing the IPs to input samples (mean values of triplicates are shown). Genome annotation is shown at the bottom according to scale with significantly up-regulated genes (adjusted p value < 0.05, Log2 fold change +/− Cu^2+^ > 1) colored in dark grey. Up-regulated genes that are bound by CopR under both conditions are highlighted with orange circles. An example of an unbound genomic region is shown in Supplementary Fig. 4A). **B**, Nucleotide enrichment analysis of upstream regions dependent on the transcription start sites comparing the sequences of selected genes (orange circles, n = 9) with the nucleotide content of sequences contributing to the consensus motif of primary transcripts in *P. furiosus.* The consensus motif, which consist of a B recognition element (BRE) and a TATA box, is highlighted above the enrichment plot (41). **C**, MEME motif analysis of selected genes. A semi-palindromic motif positioned directly upstream of the BRE element was found. **D**, Promoter sequences of genes that were further analysed by EMSA or ChIP-qPCR (compare Supplementary Fig. 4B). **E**, EMSA analysis (20 nM DNA, 0/200/400 nM protein) of selected promoter regions confirms specific binding of CopR to multiple regions, whereas no binding to a control promoter *(pf1602, gdh)* could be observed.

To elucidate the sequence specificity of CopR binding and regulation, we compared the nucleotide content of the CopR-regulated promoter regions to a background set consisting of 763 sequences that contributed to a recently published consensus motif in *P. furiosus* (41) (Fig. 4B). This archaeal-typical promoter motif is characterized by elements that facilitate transcription by the recruitment of the basal transcription factors TFB (BRE element) and TBP (TATA box) and melting of the region initially upstream of the transcription start site (TSS). In comparison to the *Pyrococcus* consensus motif, the promoter sequences of the up-regulated genes showed some minor deviations in the BRE element and the conserved A(T) at position −10 that contributes to the promoter strength (59), but the most striking difference was a C-enriched TATA box (Fig. 4B). Further upstream of the promoter sequence we identified a TC-rich and AG-rich signature from −35 to −50 that also differed from the consensus sequence content. A motif enrichment analysis using MEME identified a semi-palindromic-like motif with the minimal palindromic consensus sequence TTNNCAWWWTGNNAA, which is located at almost all CopR-regulated promoters directly upstream of the BRE element (8 of 9 with an annotated TSS) (Fig. 4C,D). Scanning of this motif in the promoter region of all known TSSs further confirmed the specificity, as all of the motif occurrences were bound and up-regulated by CopR (8 of 8 total hits, q-value < 0.01).

To validate the findings of the ChIP-seq experiments, we performed gel-shift assays and ChIP-qPCR, that both confirmed specific binding of CopR to multiple genomic regions (Fig. 4E) and enrichment under both conditions (Supplementary Fig. 4B).

Based on the genome-wide binding pattern and in combination with the results from the DGE analysis, we propose a currently undescribed regulating role of CopR on a global level to maintain copper homeostasis in *P. furiosus*.

### CopR activates transcription in vitro

To verify the stimulating role of CopR, we performed *in vitro* transcription experiments with a DNA template that allows simultaneous transcription of the divergently orientated *copR* and *copA* genes (Fig. 5A,B). In the absence of CopR, the main transcript originated from the own strong promoter, and *copA* was only weakly transcribed. However, with increasing concentrations of CopR, transcriptional output increased for the *copA* gene and *copR* transcription was significantly reduced (Fig. 5B). In contrast, it did not affect a control template, lacking the CopR binding site (Fig. 5A), which clearly indicates that CopR is responsible for both, activation of *copA* and repression of *copR.* The reason for the observed *in vitro* CopR-induced activation is not known, since the stimulating effect of *copA in vivo* was observed only under the presence of copper ions. Attempts to increase the *copA* stimulating effect by adding CuSO_4_ or AgNO3 failed. The presence of such ions inhibited the transcription in general (data not shown).

**Figure 5:**
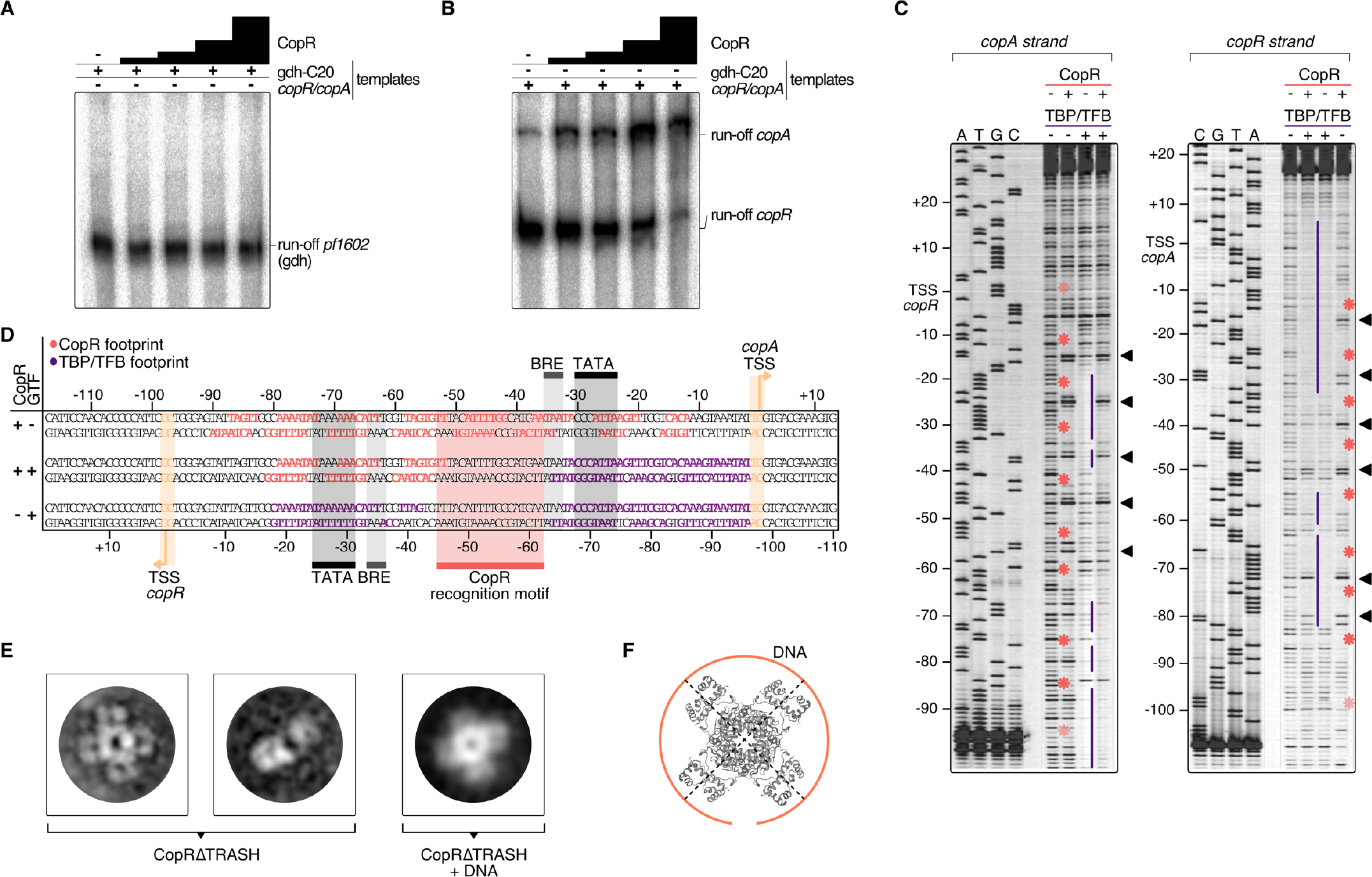
Mechanistic and structural characterisation of CopR. **A**, Influence of CopR on *in vitro* transcription. 2 nM of the *gdh* and **B**, the *copR/copA* templates were transcribed in the presence of increasing concentrations of CopR (0.3, 0.6, 1.2, 2.3 μM). **C**, DNase I footprint on the *copR/copA* template in the presence of CopR, TBP/TFB and all three components. TBP/TFB-protected regions (purple lines), CopR-protected regions (red asterisks) and hypersensitive sites (black triangles) are highlighted. **D**, Summary of the protected regions determined in the *in vitro* DNAse I footprinting assay. Promoter elements, transcript boundaries and the semi-palindromic CopR recognition motif are highlighted. **E**, Representative 2D class averages of CopRΔTRASH in the absence and presence of a *copR/copA* DNA template reveal an octameric assembly formed by a tetramer of dimers in both states. **F**, Putative model of CopR bound to DNA. The octameric cartoon structure (PDB: 1I1G) represents LrpA from *P. furiosus,* that forms a similar structure (61).

For additional information about the mechanism of activation, we performed DNase I footprinting experiments at the *copR/copA* promoter (Fig. 5C,D). The binding of CopR revealed an extended binding pattern in this region consisting of eight strong single footprints in the central region of the fragment and some additional weaker footprints towards the border of the fragment. Most of the strong signals are separated by hypersensitive sites separated by approximately one helical turn (Fig. 5C). The correlation of the binding motif obtained from the ChIP-seq experiment with the footprint pattern revealed that the motif is located between the divergent TBP/TFB binding sites of *copR* and *copA.* CopR footprint signals are positioned in the centre of the motif and nearby upstream and downstream. These signals are separated by hypersensitive sites, which touch only one or two bases of the beginning or the end of the consensus sequence. Simultaneous presence of the two basal transcription factors, TBP and TFB, and CopR in the reaction (Fig. 5D, lane 2), revealed that under these conditions only CopR is in contact with DNA in the region of the *copR* promoter. This finding is in agreement with the identified repression of *copR* in the *in vitro* transcription experiments. In contrast, the position of the CopR footprint upstream of the TBP/TFB footprint that is still present at the *copA* promoter enables CopR-mediated stimulation of *copA*.

### Towards a structural view of CopR

CopR is the crucial player in the copper-triggered differential regulation of genes that supports detoxification of the cell. At the same time, however, it is not apparent how activation is achieved considering that CopR always seems to be bound to the respective promoter regions. Therefore, we aimed to elucidate the structural properties of CopR. To this end, we developed a purification protocol for the isolation of highly pure CopR. Based on the comparison of CopR to standard calibration proteins, elution profiles of size exclusion chromatography runs (Superdex 200) indicated an octameric conformation (data not shown). As the elution profile revealed an increased symmetric peak for CopRΔTRASH in comparison to the wild type protein, we have selected CopRΔTRASH for further analysis.

Negative-stain TEM imaging confirmed an extremely high monodisperse fraction of the CopRΔTRASH mutant. Finally, 2D classification of more than 50,000 particles, gave rise to the assumption that CopR forms an octameric assembly oriented in a cruciform-like structure (Fig. 5E). Due to the high similarity of the structure to LrpA and F11 (60, 61), we assume a similar outside orientation of the helix-turn-helix domain of the four CopR dimers that positions DNA at the outside of the dimer.Furthermore, CopR complexed with DNA of the *pf706.1n* promoter did not alter the overall octameric structure of the protein (Fig. 5E). With a diameter of 16 nm, a DNA fragment of about 150 bp would be necessary to completely wrap the DNA around the protein (Fig. 5F) (61).

## Discussion

From the combination of *in vitro* based approaches, genetic manipulation and the integration of DGE and ChIP-seq data used in this study, we conclude that CopR from *P. furiosus* is a copper-sensing global regulator of transcription and essential during copper detoxification.

The importance of CopR for maintaining copper homeostasis became apparent by growth experiments. While we could show that a *P. furiosus* parental strain can adapt to μM-concentrations of Cu^2+^ (up to 100 μM were tested), the growth defect on a CopR-knockout strain was significant. We assume that limiting amounts of essential components necessary for maintaining copper homeostasis are responsible for the observed growth phenotypes. The most obvious component is the copper efflux pump CopA, which showed a 70-fold increase at the RNA level under copper-shock conditions. *In vitro* experiments confirmed that enhanced *copA* transcription is mediated by CopR-induced activation. The *substantial copA* enrichement is also in agreement with the observed CopR-binding to the upstream region of *copA* as indicated by EMSA and DNase I footprinting analysis. In summary, we assume that CopR is responsible for sensing copper concentrations and transcriptional activation of the corresponding genes necessary to maintain copper homeostasis.

Our findings for CopR are in agreement with data from *S. solfataricus* strain 98/2 and *Halobacterium salinarum* as the corresponding knockout mutants also revealed CopR as a positive regulator for *copA* transcription (29, 62, 63). In contrast, recent data from the closely related *T. onnurineus* NA1 suggested CopR as a repressor for autoregulation and *copA* transcription (28). This fundamental discrepancy for *copA* transcription is remarkable in the light of the identical organization of the operons and 79% sequence identity between both proteins instead of 30% to the *Saccharolobus* CopR. A comparison in more detail revealed that in *Pyrococcus* -in contrast to *Thermococcus-* low μM CuSO_4_ concentrations do not lead to a full release of the protein from the DNA but result in a low mobility complex indicating a conformational change of the CopR-DNA complex. These findings are also in line with *in vitro* DNA-binding studies from *S. solfataricus* strain P2, which also suggest a partial rearrangement of CopR-binding in the presence of copper instead of dissociation (25).

An additional difference between both organisms is the quaternary structure of CopR in solution: In the case of *Thermococcus* a tetrameric structure was determined using size exclusion chromatography analysis (28) and for *Pyrococcus* an octameric complex was found by negative-stain TEM imaging and gel filtration experiments. In order to explain these differences, it is tempting to speculate that the presence of a Hisβ tag in the N-terminal region of *Thermococcus* CopR is responsible for the observed different quaternary structure of the protein. Furthermore, it is possible that the presence of this tag also contributes to an increased sensitivity to copper towards dissociation from the DNA instead of allowing a conformational switch necessary for transcriptional activation. To verify the different regulation mechanism of CopR in *T. onnurineus* NA1, additional information about the susceptibility of the mentioned *copR* deletion strain (28) to increasing copper concentrations or the behaviour of CopR without a His_6_ tag would be helpful.

Despite the inconsistency of the function of CopR as repressor or activator for *copA* transcription between *Thermococcus* and *Pyrococcus*, the ChIP-seq data collected in this study demonstrate an almost identical DNA binding pattern of CopR independent of the presence or absence of copper ions. This finding also points to a required structural rearrangement of CopR on the DNA to activate transcription in the presence of copper ions. Such behaviour would be very similar to the function of the copper-sensing transcription factor CueR from *Escherichia coli* (64). CueR can activate transcription by controlling open complex formation, while it is continuously bound to DNA (65). This mechanism may permit more rapid responses to environmental changes. In an evolutionary context, the high copper-toxicity could have been a driving force for the independent development of this regulatory mechanism in different domains. Nevertheless, there is no sequence similarity between both proteins, CueR belongs to the predominant bacterial MerR family of regulators (PROSITE documentation PDOC00477), and CopR belongs to the Lrp/AsnC family (PDOC00520). The latter one is a rather old family of prokaryotic transcriptional regulators and very common in Archaea (66). Crystal structures of several archaeal Lrp members indicate a highly conserved octameric structure with an N-terminal winged HTH motif for DNA binding and a C-terminal domain necessary for oligomerization and effector binding (60, 61, 67). Our TEM imaging data of negative-stained CopR also revealed a tetrameric assembly of dimers with most likely the DNA wrapped around the protein. This finding is in line with published structures of DNA-protein complexes FL-11 and Grp (60, 67, 68) which further confirmed an accumulated occurrence of an octameric assembly within the Lrp family. The identification of extended DNA binding regions and hypersensitive sites in footprinting experiments is also in agreement with the assumed wrapping of the DNA around the octamer (69, 70). Therefore, the behaviour of CopR is very similar to published data of the Lrp family with the difference that CopR uses copper ions as effector instead of molecules of the amino acid metabolism. However, the ability to use a variety of effector molecules is pervasive for the archaeal subfamily. A detailed analysis of eight Lrp/AsnC paralogs in *Halobacterium salinarum* revealed that these proteins are involved in regulating genes in response to copper or oxidative stress, changes in K^+^ or NAD^+^ concentrations or modified growth conditions (71).

The Lrp/AsnC family is not only involved in the regulative response to a wide range of different physiological conditions but also employ different mechanisms of transcriptional regulation. Repression by preventing the recruitment of the RNAP is demonstrated for LrpA from *P. furiosus* (72) and activation by stimulating the binding of TBP is shown for Ptr2 in *Methancaldococcus jannaschii* (73). Besides, a dual regulator mechanism has been shown for Ss-LrpB from *Saccharolobus solfataricus*, which activates transcription at low factor-concentrations, whereas at high concentrations, transcription is repressed (74).

Based on our results, we conclude that CopR also has a dual function as repressor and activator, but the situation seems to differ from the LrpB from *S. solfataricus*. *In vitro* transcription experiments in the absence of CopR indicate a strong *copR* promoter without the necessity for further activation. However, *in vivo*, CopR remains always bound to the *copR* promoter region, which blocks TBP/TFB recruitment, represses transcription of its gene and only allows some basal expression. This goes in line with the DGE data, which indicate low-level expression of *copR* independent of copper ions and high-level expression of *copA* in the presence of copper. Similar results about these differences in the expression rates were also described in *Saccharolobus solfataricus* P2 (25), *Saccharolobus solfataricus* 98/2 (56), *Sulfolobus metallicus* (75), *Ferroplasma acidarmanus* Fer1 (55) and *Halobacterium salinarum* (62).

To interpret our results and to integrate these data with the knowledge gained for the Lrp family in general, we suggest the following regulation mechanism for divergent transcription (Fig. 6): Under normal growth conditions, CopR binds to multiple binding sites with about 150 bp of DNA wrapped around each octamer. Each dimer of the octamer is in “direct contact” with a weakly conserved DNA sequence as indicated by motif analysis of the ChIP-seq data and the footprinting experiments. We assume an increased affinity to binding sequences located directly upstream of promoter sequences and cooperative binding, which seems to be a common feature of Lrp molecules (76, 77), to weaker signals stimulated by the octameric structure of CopR downstream of the promoter. An additional contact of Lrp-like molecules downstream of the TATA box in combination with transcriptional activation is already described for Ptr2 and BarR (70, 78).

**Figure 6:**
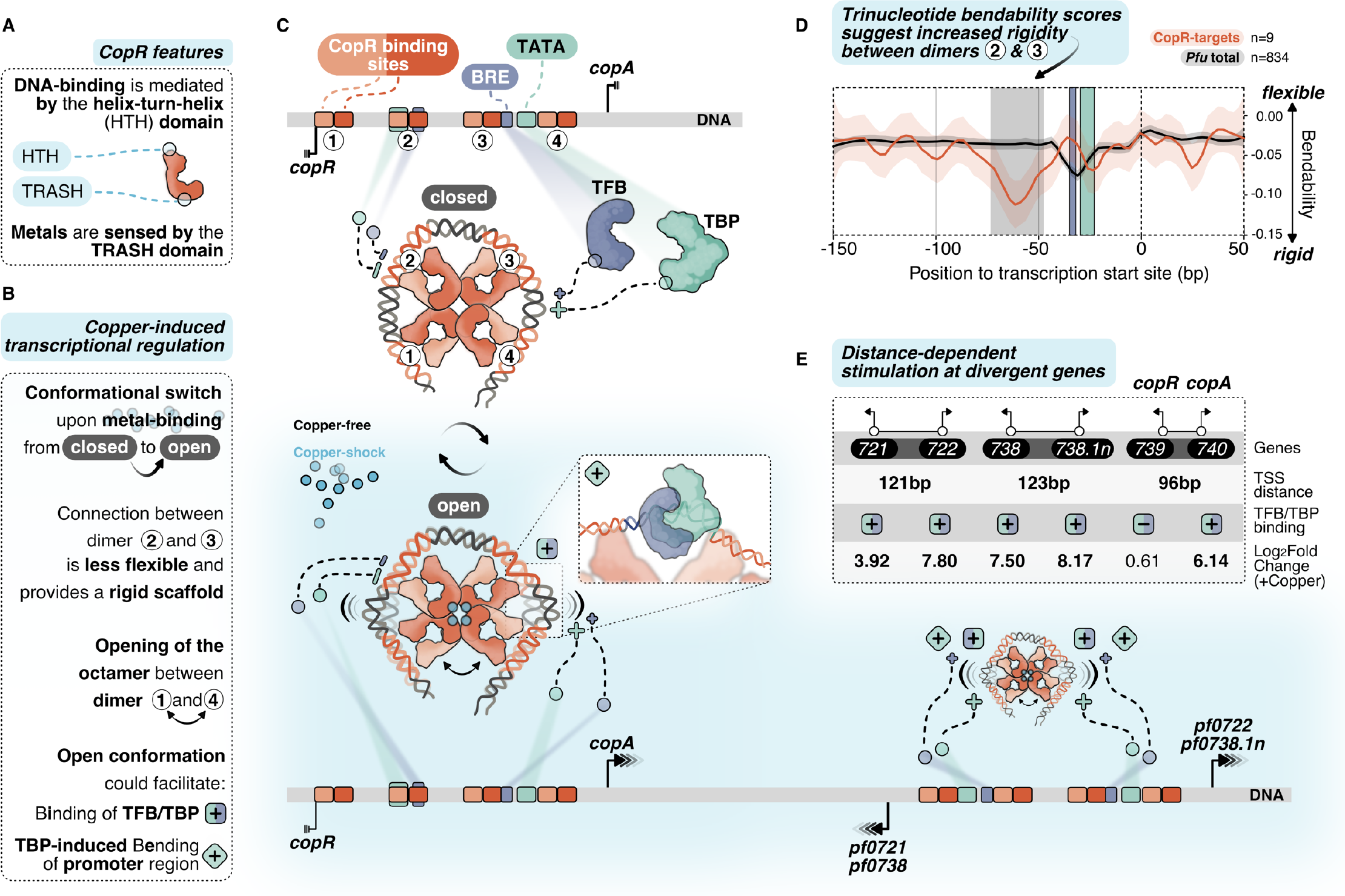
Putative model of allosteric CopR regulation in *P. furiosus.* **A**, Domain architecture of CopR (monomer shown schematically in red) highlighting the DNA-binding HTH domain and the metal-sensing TRASH domain. **B**, Binding of copper presumably triggers a conformational switch from a closed state to a complex, which is opened between dimer 1 and 4. In combination with the rigid connection between dimers 2 and 3 it is possible that this conformational switch could facilitate TBP/TFB binding (+ square) and/or TBP-induced bending to the corresponding promoter regions (+ turned square). **C**, CopR-regulated promoter regions include CopR-binding sites (orange/red) and the archaeal-specific promoter elements BRE (recruits TFB, purple) and the TATA box (bound by TBP, light-green). Transcription start sites (TSS) are indicated by vertical lines. Transcription is either repressed (3 lines) or stimulated (arrows) under copper-shock conditions (lower panel, light-blue). The octameric CopR assembly in open conformation allows bending of critical promoter regions and facilitates binding of TBP/TFB depending on the distance of the divergent TSS: While CopR prevents binding of general transcription factors to the *copR* promoter, TBP/TFB can bind to the second promoter (*copA*). Simultaneous binding of CopR and two sets of GTFs stimulates transcription in both directions for *pf0722/pf0271* and *pf0738/pf0738.1n.* **D**, Major groove bendability of selected promoters revealed increased rigidity between dimers 2 and 3. Major groove bendability of promoter sequences was estimated based on trinucleotide scales derived from DNase-I cutting frequencies (80, 81). More negative values are the result of less cutting by DNase-I and indicate that the DNA is not bend towards the major groove and is therefore less flexible. Trinucleotides were extracted from CopR-target promoters (n=9) and all available promoters defined in *P. furiosus* previously (41) from −150 bp to +50 from the TSS. Each line represents the smoothed conditional mean with confidence intervals (0.95) displayed as shaded areas. Boxes represent area between dimers 2 and 3 (grey), BRE (purple) and TATA box (light-green). **E**, Summary of the TSS-distance dependent stimulation of divergent CopR-regulated transcripts.

For transcriptional activation, we suggest an allosteric regulation mechanism where the binding of effector molecules (most likely Cu^+^) is sensed by the TRASH domain alone or in combination with the HIS stretch at the C-terminal end (Fig. 6A). The role of the TRASH domain in copper binding is indicated by the decreased metal sensitivity of the ΔTRASH mutant in the gel shift assays (Supplementary Fig. 2D) and was also previously demonstrated by mutational analysis in *T. onnurineus* NA1 (28). Due to the binding of the metal, we expect a conformational switch resulting in the opening of the quaternary structure of the octamer similar to the structure of Lrp from *Escherichia coli* or FL11 with bound arginine from *Pyrococcus* OT3 (68, 79). We assume that opening between dimer 1 and 4 is preferred due to increased flexibility at this position as these dimers are not directly linked with the wrapped DNA (Fig. 6B,C). In contrast, there is a direct connection from dimer 1 to dimer 2, 3 and 4, which most likely provides a more rigid scaffold. The bendability of CopR-regulated promoters in comparison to the total set of promoters in *P. furiosus* (41) was estimated by comparing trinucleotide scores (80, 81) and clearly shows a less flexible region between dimer 2 and 3 (Fig. 6D). The movement of dimer 1 towards 2 and dimer 4 towards 3, initiated by the opening of the octamer, may reduce torsional stress on the DNA between these corresponding pairs of dimers, which could either facilitate the accessibility of TBP and TFB to the corresponding promoter sequences or enable TBP-induced bendability or is involved in both (Fig. 6B,C). Torsional stress on the DNA as limitation for binding of TBP was already demonstrated by single molecule FRET experiments in *Methanocaldococcus jannaschii* (82).

Interestingly, provided a divergent gene organization and an appropriate distance between the two TSSs, one CopR octamer can promote transcriptional activation of two separate gene clusters (Fig. 6E). Simultaneous stimulation by CopR requires a distance of about 122 bp between the two TSSs *(pf0721/pf0722; pf0738/pf0738.*1n). In contrast, the reduced distance of 96 bp in the case of the *copR/copA* gene cluster leads to repression of the *copR* gene independent of the presence of copper ions, most likely due to binding interference of dimer 3 with the *copR* promoter. Additionally, there is also the possibility of activation at only one side of the octamer, whereas the position of the other side is located at the end of another gene *(pf0726/pf0727).*

Besides mechanistic details of CopR transcriptional regulation, our data also allow insights into a general copper-specific transcriptomic response (Fig. 7). To avoid a transcriptomic response due to cell-death rather than a metal-specific response, we applied a moderate copper shock using 20 μM CuSO_4_ in all *in vivo* experiments. However, under these conditions, we already saw a strong phenotype of the CopR knockout mutant, which emphasizes the essential role of CopR. Using this “semi-toxic” concentration in the DGE analysis, we found 34 strongly up-regulated genes (> 2-fold) upon copper-shock (Supplementary Table 3). The transcriptional pattern under copper shock conditions is comparable to other metal stress transcriptomic responses in prokaryotes and eukaryotes, and apart from metal-specific genes also includes non-metal related genes that cooperatively contribute to metal resistance (12, 83).

**Figure 7:**
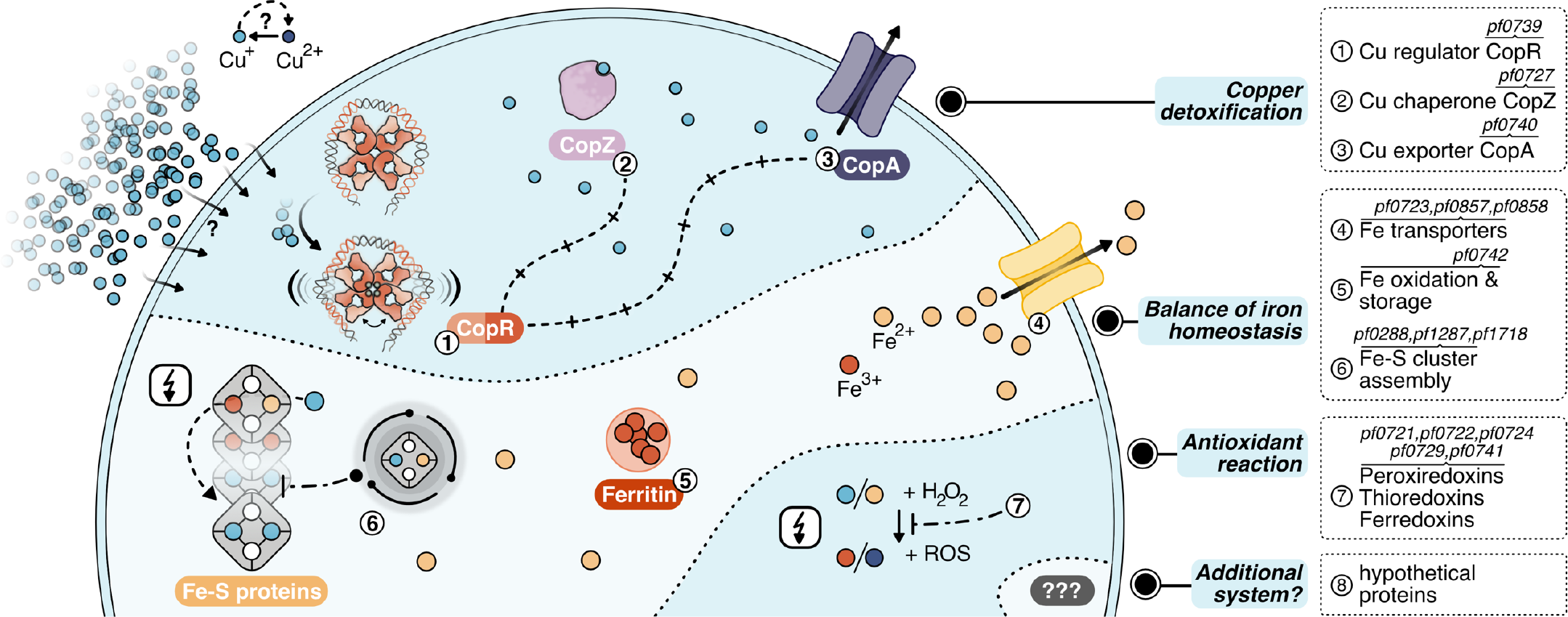
Layers of copper detoxification in *P. furiosus*: In a primary response, CopR (1,orange-red) senses excessive amounts of Cu^+^ ions (lightblue), that entered the cell by a currently unknown mechanism or diffusion and activates the transcription of the copper-chaperone CopZ (2,light-purple) and the copper-exporter CopA (3,purple). After CopZ potentially delivers the Cu^+^ ions to the transporter, they are exported out of the cell. The main toxic effect of copper ions is caused by replacing iron ions in iron-sulfur clusters. Therefore, the induction of Fe-S cluster assembly proteins, Fe-transporters and ferritin helps to re-balance iron homeostasis (4–6). Additionally, antioxidant enzymes prevent the induction of the Fenton reaction (7), which otherwise causes toxicity by the production of reactive oxidative species (ROS).

For the primary detoxification mechanism, *Pyrococcus* relies on the induction of the copper efflux pump CopA and the metallochaperone CopZ (PF0727), which directly interact with the copper ions. An additional cluster of genes deals with i) iron homeostasis involving several transporters to pump iron ions (PF0723, PF0857, PF0858) (Zhu et al., 2013), ii) ferritin (PF0742) which combines oxidation of Fe^2+^ to Fe^3+^ together with storage of the oxidized iron inside the protein cavity (87) and iii) Fe-S cluster assembly proteins (PF0288, PF1286, PF1287, PF1718). This collection of genes fit well into the recently emerging concept that the primary toxic effect of copper is the replacement of iron in iron-sulfur cluster proteins and not the conversion of H2O2 to hydroxyl radicals (15, 83). Therefore, the induction of these proteins helps to re-balance displaced iron ions and to avoid inactivation of iron-sulfur proteins. Besides, antioxidant enzymes as peroxiredoxins, thioredoxins or ferredoxins (PF0721, PF0722, PF0724, PF0729, PF0741) can also assist in preventing the induction of the Fenton reaction by the released Fe^2+^ or Cu^+^ ions. This is in line with the finding that some of these enzymes are also induced after exposure to hydrogen peroxide (Strand et al., 2010). Since the constitutively expressed superoxide reductase also produces hydrogen peroxide (86, 88), it is most likely that the induction of these antioxidant enzymes successfully inhibits the production of hydroxyl radicals via the Fenton reaction under this low dose of copper. In consequence, there is almost no induction of genes dealing with general stress response or DNA repair mechanisms. In contrast, a copper shock in *Metallosphaera sedula* induced a mixed gene population of metal-specific and also generic responses indicating that the conditions used have had much more potent effects concerning viability in comparison to our setup (89).

In this context, it is interesting to note that *Metallosphaera* uses an additional mechanism for copper resistance: sequestration with inorganic polyphosphate to facilitate export from the cytoplasm (90). Such a mechanism is also described for *S. solfataricus*, as a mutant strain-unable to accumulate polyphosphate-showed an increased copper sensitivity in spite of *copA* up-regulation (91). Based on our data, there is no indication that a comparable system is implemented in *Pyrococcus.* Nevertheless, the induction of eight hypothetical proteins opens up the possibility that different sequestration systems or an additional mechanism for copper detoxification exist.

## Data availability

The R scripts detailing the analysis can be found in the corresponding Github repository under www.github.com/felixgrunberger/CopR.

Raw sequence data have been uploaded to the SRA and are available under project accession number PRJNA603674.

## Acknowledgements

The authors thank Renate Richau and Wolfgang Forster for excellent technical assistance, Dominik Strobel for constructing a CopR expression clone, Markus Schick and Christine Lindenthal for initial work in the setup of the genetic system and Thomas Kopp from the electronic workshop from the University of Regensburg for the setup of the optical device to measure cell density.

## Author contributions

FG did the DGE and the complete bioinformatic analysis, RR constructed the *Pyrococcus* deletion strain, IW did the *in vitro* transcription and the footprinting experiments. VN and LK performed the gel shift assays, KB the ChIP-seq experiments and the qPCR assays. MK, NW, ZE, MGM and CZ did the negative stain imaging. FG, DG and WH wrote the manuscript and DG and WH coordinated and supervised the work. All authors agreed to the final version of the manuscript.

