## Supplemental Figures for "CopR, a global regulator of transcription to maintain copper homeostasis in *Pyrococcus furiosus*"

**SUPPLEMENTARY FIGURES**

### Supplementary Figure 1

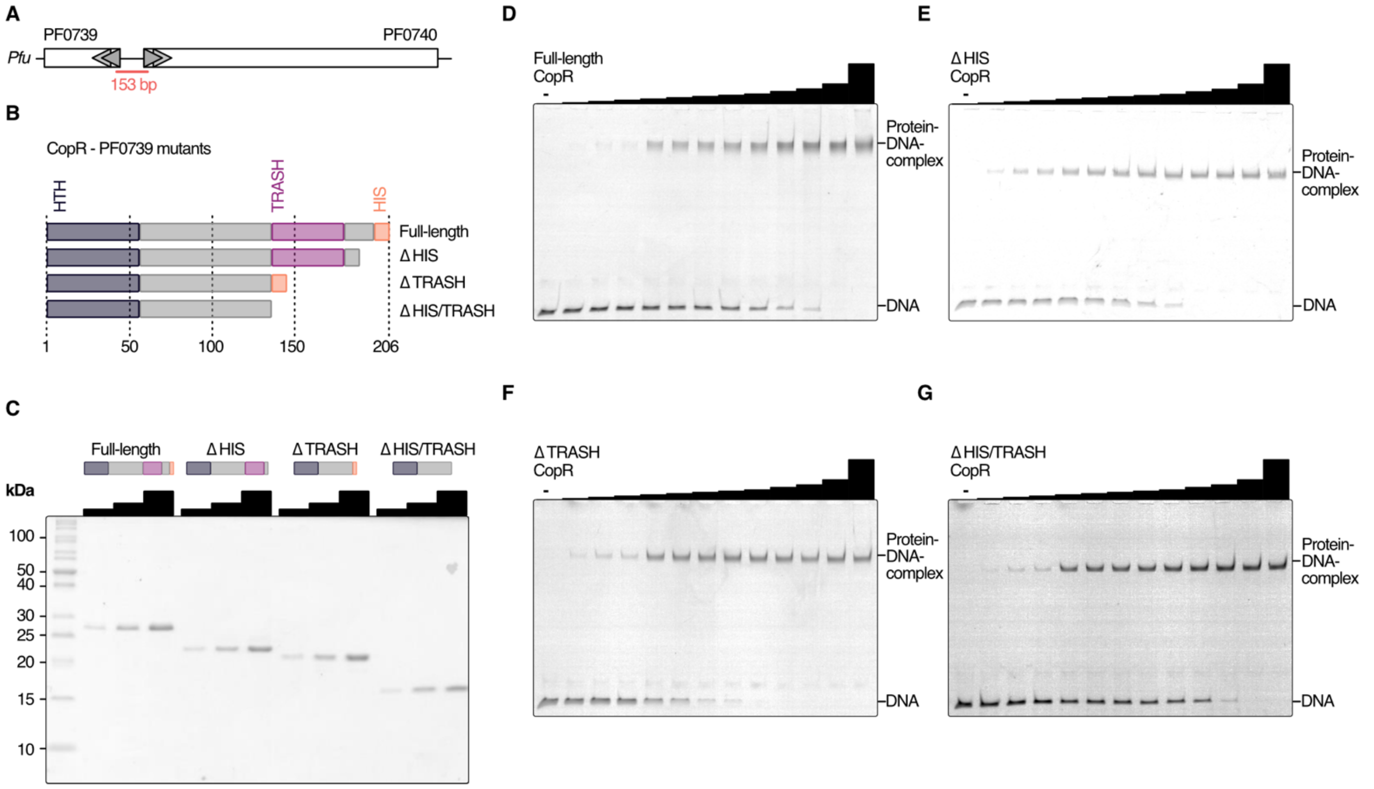


**Supplementary Figure 1. CopR (PF0739) binds to the promoter region of *copA* (PF0740). A**, Schematic representation of the *copR/copA* locus in *P. furiosus*. The template used for the EMSA analysis is highlighted in red and contains both translation start sites. **B**, Schematic of mutants generated for the functional characterisation of CopR. DNA-binding helix-turn-helix (HTH), metal-sensing TRASH domain and additional C-terminal Histidine-rich sequence are highlighted in different colors. **C**, SDS-PAGE analysis of purified recombinant proteins. **D**, EMSA analysis were performed using 20 nM of DNA (153 b, see panel a) and increasing concentrations of recombinant protein (12.5, 25, 37.5, 50, 62.5, 75, 87.5, 100, 125, 150, 200, 400 nM).

### Supplementary Figure 2

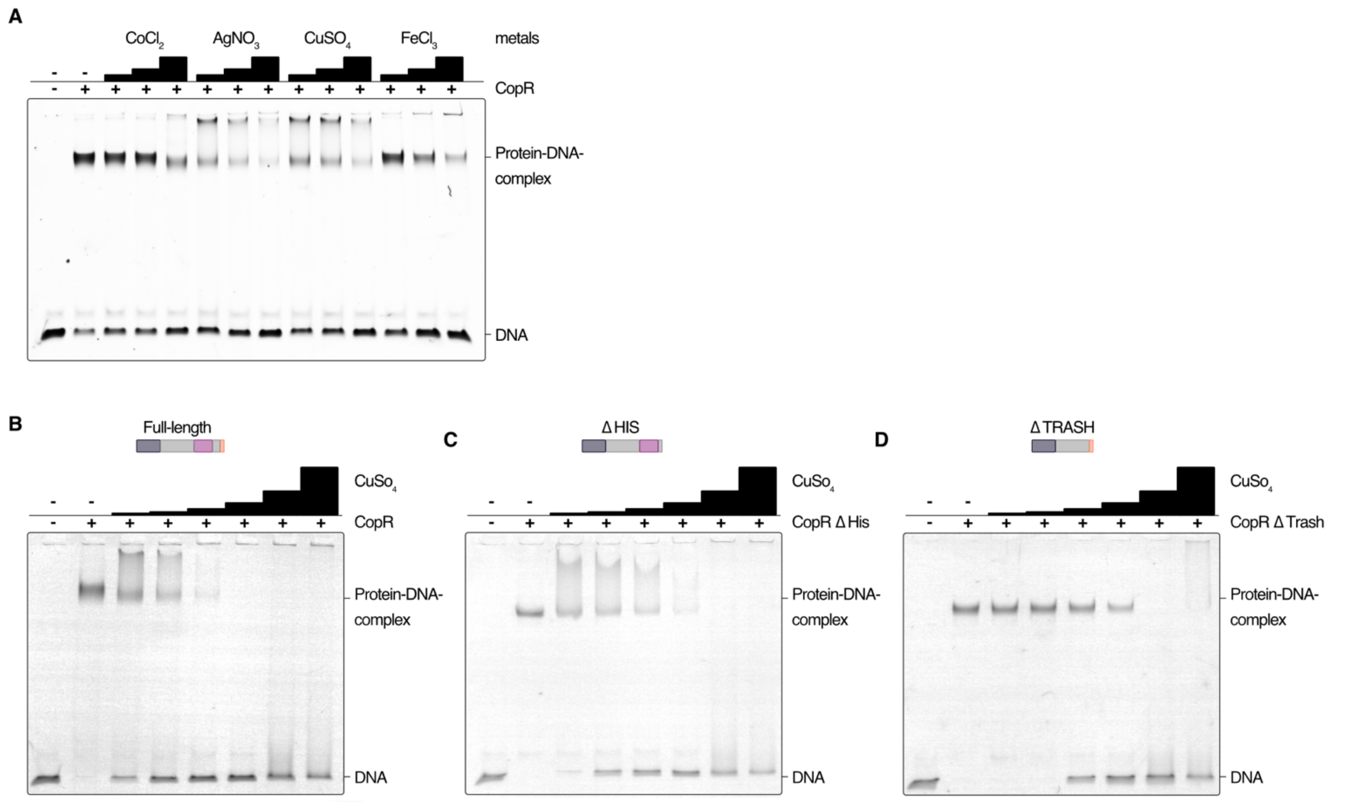


**Supplementary Figure 2. Metal specificity of CopR and domain-deleted mutants in *P. furiosus*. A**, EMSA analysis was performed using 20 nM DNA (*copR/copA* promoter), 200 nM full-length protein and increasing concentrations (12.5, 25, 50 µM) of the respective metal ($CoCl_2$, $AgNO_3$, $CuSO_4$, $FeCl_3$). **B**, Influence of increasing $CuSO_4$ concentrations (50, 100, 200, 400, 800, 1600 µM) on the DNA-binding behavior of full-length CopR, **C**, CopRΔHIS and **D**, CopRΔTRASH using 20 nM DNA and 200 nM protein.

### Supplementary Figure 3

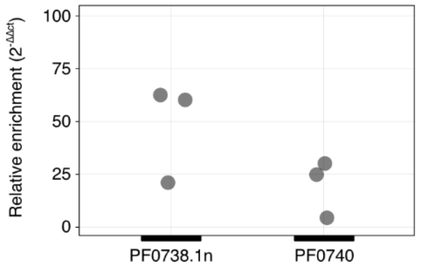


**Supplementary Figure 3. Relative enrichment of *pf0740* and *pf0738.1n* measured by RT-qPCR.** Expression levels from biological triplicates (individual points are shown) were compared to a house-keeping gene *pf0256*. Enrichment was calculated using ΔΔct method.

### Supplementary Figure 4

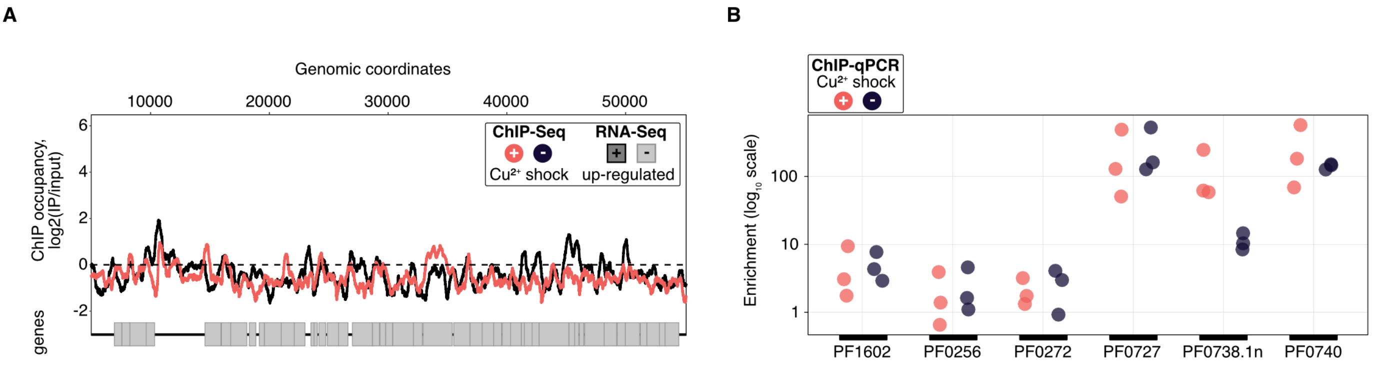


**Supplementary Figure 4. Confirmatory analysis of ChIP-seq results. A**, Exemplary CopR-unbound 50-kb region of *P. furiosus* (compare Fig. 4A). ChIP-seq curves were generated for $Cu^{2+}$ shocked (red) and untreated (dark blue) samples by comparing the IPs to input samples (mean values of triplicates are shown). Genome annotation is shown at the bottom according to scale with significantly up-regulated genes (adjusted p value < 0.05, $Log_2$ fold change +/- $Cu^{2+}$ > 1) colored in dark grey. **B**, ChIP-qPCR results of multiple ChIP-seq identified CopR-unbound (PF1602, PF0256, PF0272) and CopR-bound (PF0727, PF0738.1n, PF0740) regions. Data are shown as individual points of biological replicates for normal conditions (dark blue) and copper-treated cells (red). Values are calculated as fold enrichment over input sample.
